# ‘PePApipe’: a complete bioinformatics analysis pipeline for African Swine Fever Virus genome

**DOI:** 10.64898/2026.01.12.698977

**Authors:** Vicente Lopez-Chavarrias, Irene Aldea, Jovita Fernández-Pinero

**Affiliations:** Centro de Investigación en Sanidad Animal (CISA), Instituto Nacional de Investigación y Tecnología Agraria y Alimentaria (INIA), Consejo Superior de Investigaciones Científicas (CSIC), Valdeolmos, Madrid, Spain

## Abstract

African Swine Fever Virus (ASFV) is of high concern in porcine livestock across the world due to both the high mortality rates and the trade restrictions imposed on affected regions. Viral genome is large and complex, but genomic analysis is essential for tracing its origin and evolution. Although several bioinformatics tools exist for genome assembly and analysis, no single platform integrates all necessary steps in an accessible and systematic way. In this study the authors developed ‘PePApip’, a custom-built, user-friendly pipeline that enables rapid, complete, and efficient ASFV genome analysis. It is specifically designed for laboratory professionals with limited bioinformatics experience, requiring only basic command-line knowledge.

Starting from raw sequencing data, ‘PePApipe’ integrates thirteen software tools into one automated workflow, covering quality control and pre-processing of raw reads, *denovo* genome assembly and variant calling. Programmed in Phyton, it can be executed locally through bash scripts, or using a SLURM protocol for batch processing of multiple samples. The main outputs are the ASFV consensus genome sequence and a file listing its putative variants compared to the selected reference genome.

PePApipe classifies generated files into structures folders and produces intermediate files that can be used as inputs for further or parallel analyses; also users can enable or disable specific steps in each particular case. This pipeline is adaptable and complementary to downstream steps such as viral genome annotation or genome visualization.

By consolidating all stages of viral genome analysis into a single automated workflow, PePApipe reduces the likelihood of user error and enhances reproducibility and efficiency. This easy-to-use pipeline will facilitate the transition from sequencing to assembly and analysis of viral genomes, ensuring a fast and reliable response to molecular analysis demands. Finally, the pipeline can be easily adapted to the study of other viral species, expanding its application in infectious diseases surveillance.

**Author summary:** African swine fever is a devastating viral disease that threatens pig production worldwide, causing severe economic losses and limiting international trade. Tracing and understanding how the virus spreads and evolves are key for preparedness and control of the disease, currently being a major sanitary challenge. However, viral genome analysis is often technically demanding and difficult to standardize, especially for laboratories without specialized bioinformatics expertise. In this study, we present PePApipe, an automated and user-friendly computational pipeline designed to simplify and standardize the complete genomic analysis of African swine fever virus. Starting directly from raw sequencing data, PePApipe guides users through all essential steps, from quality control to genome assembly and variant detection, producing reliable and reproducible results in a short time. By integrating multiple established tools into a single workflow and providing clear intermediate outputs, our approach reduces the risk of user error and increases transparency. Importantly, we designed PePApipe with accessibility in mind, enabling laboratory scientists to perform advanced genomic analyses with minimal computational background. While developed for African swine fever virus, the pipeline can be adapted to other viruses, making it a flexible resource for viral genomics, outbreak investigation, and future data-driven research.

## Introduction

Despite the availability of generic bioinformatics tools for genome analysis across a variety of organisms, the intrinsic biological and structural differences between genomes often necessitate the development of specialized tools tailored to the organism of interest—this is particularly true in the case of viruses. One such organism is the African swine fever virus (ASFV), a large, complex, double-stranded DNA virus belonging to the family *Asfarviridae* (1). ASFV is of critical concern to the global swine industry due to its ability to cause a haemorrhagic disease with near 100% mortality in domestic pigs, as well as the substantial trade restrictions imposed on affected regions following confirmed outbreaks.

The absence of an effective vaccine, coupled with the virus’s complex biology and high environmental stability, makes ASFV a formidable challenge for veterinary and agricultural systems worldwide (2, 3). Indeed, ASFV has garnered increasing scientific interest in recent years due to its severe threat to global pig production and food security derived from the worldwide epidemiological scenario. The virus has expanded beyond its original endemic regions in sub-Saharan Africa, initially entering Eastern Europe in 2007 and further spreading throughout Europe, Asia, and reaching the Americas in 2021, which has further intensified the need to understand its pathogenesis, transmission mechanisms, and evolutionary dynamics (4, 5). Moreover, knowing the origin of a new ASFV incursion and tracing the virus spread are critically demanded right after an outbreak occurs in a new region. Besides, the socioeconomic consequences of ASF outbreaks, particularly in countries with major swine industries such as China or Germany, have highlighted the urgent demand for targeted control strategies and improved diagnostic tools (6, 7).

The analysis of the ASFV genetic content is key to infer the possible origin of the viral incursion and to inform possible measures that can be put in place to avoid further transmission of the virus, thus enabling a full epidemiological picture of the situation which in turn can help bring the situation back under control.

The problem with viruses in general, and with the ASFV in particular, is that their genome is prone to change (8), and this variability in nature is typically a consequence of its replication *in vivo* when it infects host cells (domestic pigs and wild boars). With a genome of variable size between 170 to 194 kbp, ASFV is able to exchange large parts of its DNA with itself or with variants of the same virus in the cell environment of the host, with the resulting consequence of producing new genomes. Genome variations can be simply point mutations or small indels (insertions and deletions) occurring not very frequently, but nevertheless altering the original genomes, or these changes can be large indels leading to gene rearrangement, and homologous recombinations could occur as well (9). An additional peculiarity of ASFV genomes is that both left and right terminus regions of its DNA are very variable compared to the central section; moreover, these end regions contain inverted terminal repeats which further complicate both the assembly and annotation of the genomes at the analysis stage (10).

Although several tools exist that can be used to perform assembly and analysis of DNA genomes of viruses (https://zenodo.org/records/7764938), such as ASFV (11, 12), the authors of this work have not found to date a tool able to compile all steps involved in this analysis in a simple and accessible way.

The present tool, called ‘PePApipe’, is fit for this purpose, and implements the work in a rapid, complete, efficient and clear manner. Moreover, this pipeline has been developed bearing in mind that end-users are mostly laboratory professionals with limited bioinformatics skills.

## Results and discussion

The pipeline described on this paper has been built having into account the specific characteristics of ASF viruses. Every step was tried on the sequences from the six samples using all tools known to the authors to be adequate for that purpose. After different runs, the results using different tools were compared with the rest and the best among them were used to decide upon which tool to include in the pipeline.

To examine the quality of raw reads we included FastQC (13) only once for an initial quality assessment and used fastp (14) instead for the post-trimming quality assessment because fastp includes this assessment as a subsequent step to the trimming process. To decide on a trimming tool we considered trimmomatic (15), cuatadapt (16) and fastp (14), and we opted for the latter because of the post-trimming quality check that this tool provides, being the average length of the resulting fragments similar with the three tools (data not shown).

We tested the trimming with the three tools mentioned applying a common minimum length for trimmed reads of 60 and two minimum quality thresholds: 20 and 30. The reads obtained with quality thresholds of 20 were consistently longer than the reads obtained when we were stricter with the quality threshold (for example 30), hence we opted to include quality thresholds of 20 in the trimming tool. In our experience, when working with viruses, unless the analyst has an exceptionally good sequencing sample, it is better not to be too strict on quality of reads, particularly when the amount of viral DNA at the outset of the experiment is not too high, even after using viral enrichment methods to prepare specific libraries prior to sequencing, as it was our case.

For the *de novo* assembly stage we focused on the tool Unicycler (17) which has been shown to perform well on assembly of viruses. Moreover, although it is a tool designed to combine short and long reads to perform a hybrid assembly, it delivers similar results with viruses working with short reads only, which is an advantage in this case. Of course, the tool is equally capable of analysing a mixed of viral short and long reads. Unicycler already includes other assembly and polishing tools such as Spades (18) and Pilon (19).

Two different software programmes were tested for *de novo* assembly of the sequences: SPAdes and Unicycler (Table 1). The Unicycler software was used with the parameter --linear_seqs 1, forcing it to return a single contig. We observed a significant difference in the number of contigs corresponding to ASFV obtained with each programme. The number of contigs obtained with SPAdes ranged between 16 and 504 whereas with Unicycler was between 3 and 6, demonstrating the extra scaffolding gained with other programmes included in Unicycler such as Pilon (19).

**Table 1.** Comparison of parameters obtained using SPAdes and Unicycler as de novo assembly tools for ASFV sequences.

| SAMPLE | ASFv contigs<br>SPAdes | ASFv contigs<br>Unicycler | Larger contig<br>SPAdes | Larger contig<br>Unicycler |
| --- | --- | --- | --- | --- |
| E1 | 16 | 3 | 185354 | 185352 |
| E2 | 120 | 3 | 185506 | 185352 |
| E3 | 120 | 4 | 115053 | 185352 |
| E4 | 256 | 6 | 113089 | 185352 |
| E5 | 504 | 3 | 30747 | 125185 |
| E6 | 117 | 4 | 145211 | 185353 |

It is worth noting that we obtained more contigs with SPAdes than those shown in Table 1, but we have only displayed the contigs corresponding to the virus of interest. In Sample E2, for example, nearly 9000 contigs were obtained that did not correspond to ASFV, but to other organisms. This large number of contigs obtained with SPAdes complicates the analysis of the virus that is of interest to us.

Differences in the length of the longest contig were also observed. With Unicycler, we obtained a contig between 40k and 90k nucleotides longer in 4 out of the 6 samples. In the samples where we obtained shorter contigs with Unicycler, the difference ranged from 2 to 154 nucleotides, therefore we chose Unicycler as the best *de novo* assembling tool for the virus of interest.

One of the main challenges when performing bioinformatics analyses of viral sequences is the need to obtain the truest and most accurate consensus sequence of the virus being analysed. Any rough approximation is useless and we must aim to obtain the true genetic picture of the virus we are confronting to. To attain this goal, in our pipeline we used a combination of two published and reliable analysis tools: RagTag and MeDuSa.

Two of the four available functionalities from RagTag (20) are used in the pipeline: RagTag Correct and RagTag Scaffold. Most of the contigs are joined together using a virus reference genome as a guide but without inclusion of any of the nucleotides from the viral reference sequence. Once this compilation job is done, MeDuSa (21) implements a final join of the remaining contigs, this time using as references as many as we can/want to provide. In our case we used 10 references of ASFV genotype II, as these were phylogenetically closest to our problem viruses.

MeDuSa attempts at further connect contigs by matching our problem virus with the sequences of the references provided, and similarly to RagTag, it does not use any of the nucleotides off the references themselves.

In our experience, approximately 80% of the assemblies we obtained were reduced to a single contig, and in the remaining 20% of cases the final contig sets included up to three, and in very few occasions, up to five contigs. The two main reasons for not being able to assemble to a final single contig were either lower quality of reads or possible contamination (in our case with bacterial plasmids of around 5,000 nucleotides).

For the alignment with reference (mapping) of our sequences we tested mainly two viral references: ASFV Georgia 2007/1 (GenBank assembly GCA_003047755.2) and ASFV Lithuania LT14/1490 (GenBank assembly GCA_008932025.1). Although all results were very similar in both cases in terms of quality and metrics, we opted for ASFV Georgia 2007/1 for all the analyses, as it represents the initial reference strain of the entrance of ASFV genotype II into the Eurasian territory

For the mapping, three different software programmes were tested: minimap2 (22), Bowtie2 (23) and bwa-mem2 (24) (Figure 2). Bowtie2 was used with two different parameter sets: --very-sensitive-local and the default parameters. Both minimap2 and bwa-mem2 perform local alignment by default, meaning that the entire read does not need to match the reference genome. Bowtie2 does not use this type of alignment by default, so the corresponding parameter must be added manually.

The percentage of reads mapping to the reference was higher in all cases with the bwa-mem2 software. As expected, the lowest percentages were obtained with Bowtie2 using the default parameters, although in five out of the six samples analyzed, Bowtie2 with the local mapping parameter was the second software that mapped most reads (Figure 2). The bwa-mem2 software was ultimately selected because it achieved higher percentages of mapped reads in all samples, optimizing the variant calling process.

This pipeline yields optimal results when used in conjunction with an enrichment strategy designed to isolate and characterize viral sequences, such as through a custom-built viral genome library or by using amplicon-based approach. In the absence of such a capture step, the proportion of non-viral reads is likely to be very high, which can significantly reduce the reliability of the results compared to those obtained with prior enrichment.

Table 2 shows, as an example, a summary of results and statistics of complete genome analyses from six different ASFV viral samples (in columns). The different metrics shown (in rows) give an idea of the variability observed among these viral samples.

**Table 2.** Example of summary of results and statistics of complete genome analyses of several viral strains.

| Statistics | Sample E1 | Sample E2 | Sample E3 | Sample E4 | Sample E5 | Sample E6 |
| --- | --- | --- | --- | --- | --- | --- |
| Per base<br>sequence<br>quality<br>(average<br>R1+R2) | 36 | 36 | 36 | 36 | 36 | 36 |
| % G+C<br>(average<br>R1+R2) | 39% | 38.5% | 39% | 39% | 41% | 38% |
| Total number<br>of reads | 1,049,376 | 1,211,198 | 1,298,236 | 1,486,248 | 2,055,450 | 1,282,808 |
| Duplication<br>rate | 33.9% | 17.4% | 18.7% | 22.3% | 18.5% | 15.7% |
| Reads passing<br>quality filters | 1,038,682 | 1,198,676 | 1,283,168 | 1,461,022 | 2,039,472 | 1,268,262 |
| % reads<br>passing<br>quality filters | 99.98% | 98.97% | 98.84% | 98.30% | 99.22% | 98.87% |
| Reads<br>mapped to<br>Reference<br>genome | 1,036,938 | 1,009,847 | 976,483 | 1,142,610 | 1,897,109 | 1,141,496 |
| % reads<br>mapped to<br>Reference<br>genome | 99.83% | 84.25% | 76.10% | 78.21% | 93.02% | 90% |
| Mean coverage per contig (depth or vertical coverage) | 795 | 777 | 751 | 863 | 1,260 | 877 |
| No. of initial fragments before MeDuSa step at *de novo* assembly | 2 | 3 | 2 | 3 | 5 | 2 |
| No. of final fragments after MeDuSa step at *de novo* assembly | 1 | 1 | 1 | 1 | 1 | 1 |
| Total *de novo* assembled length (horizontal coverage) | 187,794 | 187,892 | 187,891 | 187,898 | 188,341 | 187,893 |

The analysis of variants in genomic sequences that are subject to high variation, such as those obtained with viruses, is not an easy task. ASFV contains a DNA genome and is expected to show a low level of mutation. However, its large size (up to 194 kbp) and its ability to delete, insert, duplicate, invert, and recombine short and large genome regions, make variant calling a challenging step. We explored several tools to try to standardize this search. First we used the Nextflow platform (25) nf-core/viralrecon (26), which is a tool that was developed during the Covid-19 pandemic to analyze new coronavirus variants. We found the installation and processing cumbersome, and the outputs did not fulfil the expectations we had beforehand on this tool.

After searching for other appropriate tools we decided to compare two bayesian inference searching tools (BCFtools (27, 28) and Freebayes (29)) with a non-bayesian tool (VarScan 2) (30), using an additional fourth tool (basic variant detection included in the CLC Genomics Workbench software) (31) to analyze the overall results from four perspectives, performing an orthogonal analysis. The main finding after using the four tools was the lack of variants found by the two Bayesian methods, compared to VarScan 2 which found several variants. Overall, CLC found the same variants as VarScan 2, and the minimal discrepancies between the two were mostly related to considering or not considering non-canonical nucleotides as variants.

In our opinion, it is hard to believe that the viral sequences we analysed did not have any variants with regards to the references used, considering the high variability observed in viruses. Therefore, BCFtools or Freebayes could be more adequate for other types of genomes, and we decided to opt for VarScan 2 as the main tool for variant calling.

The validation of variants provided by VarScan 2 was carried out using a visualization tool [IGV (Integrative Genomics Viewer)] (32–35), which is the software tool most universally used for visualizing genome mappings. This mapping is an optional step and is not an integral part of the pipeline.

The *de novo* consensus sequence built by the pipeline can be annotated using a reference genome of choice. Although this is neither an essential step in the pipeline but rather a complementary analysis that can be carried out alongside, it is an essential task to identify and tag the different genes that have been assembled together. A popular tool released so far for this purpose is a universal tool for general annotation that works with viruses, and specifically with ASFV, called GATU (General Annotation Transfer Utility) (36).

To our knowledge, two other pipelines have been recently described to specifically analyse ASFV genome reads, revealing the necessity and relevance of ASFV tailored bioinformatics solutions. The first, published by Spinard *et al.* in August 2024 (11), is designed to process both short and long reads. The second, ANASFV (37), developed by Li *et al.* and released in September 2024, is specifically tailored for long-reads.

The pipeline developed by Spinard *et al.* was tested on two genomes only, while PePApipe was evaluated with six genomes. Although expanding the number of samples for testing would be ideal, this is not always feasible. Nevertheless, additional samples can be obtained from public repositories to increase the dataset size; however, the sequencing quality or the total viral read count from such sources may vary and not always meet desired standards.

For short-read analysis, Spinard *et al.* employed a reference-based approach, which, as mentioned earlier, can introduce identification bias by limiting the discovery of novel genomes. On the other hand, their use of *de novo* assembly with both short and long reads helps resolve or better elucidate inverted terminal repeat (ITR) regions compared to short reads alone. To address this limitation, PePApipe also supports a combined approach using short and long reads, with long reads incorporated in the initial *de novo* assembly step using Unicycler. One important difference between the pipeline by Spinard *et al.* and PePApipe is that the former allows submission of up to four datasets for analysis, whereas PePApipe can process as many samples as needed in batch mode, albeit at the expense of longer processing times.

In our analyses more than 50% of Illumina reads mapped to both the ASFV virus and *Sus scrofa* (host) genomes. The straight removal of all reads mapping to *Sus scrofa* would mean losing as well an important amount of reads mapping to the ASFV virus, which are essential for the viral genome assembly. Moreover, the pipeline by Spinard *et al.* does not use reads mapping to both the ASFV virus and host to find the most similar reference to their genome (among 33 reference genomes), and this may result in an inadequate identification of such reference genome. Although 10 reference genomes were used in PePApipe via MeDuSa, a larger panel of reference sequences may be used. The final consensus sequence may be more reliable at the expense of prolonging processing times, although the assembly improvement is likely to be marginal.

The first pipeline by Spinard *et al.* uses the *de novo* assembled contigs to predict a closely related reference genome whereas PePApipe do not make inferences about a reference genome but constructs the *de novo* assembly in one go by combining the use of RagTag and MeDuSa.

Moreover, Spinard *et al.* use reference genomes to align ITR segments that remain unaligned and flanking those ITR segments that could be aligned in the first place. This practice can curtail the possibilities of correct alignment of ORFs not present in such reference genomes but present in a potential ‘new’ virus.

The search for variants strategy of Spinard *et al.* is based on the use of BCFtools but we hardly found variants using this tool, as explained above. The reason for this difference could be that we used samples that were very similar between each other and between them and the reference genome sequence, as they were all ‘in vivo’ passes of the same virus. Using BCFtools with a more heterogeneous dataset may result in more variants called.

The publication describing the ANASFV pipeline does not provide as detailed an explanation of the assembly stage as PePApipe or the pipeline by Spinard *et al.* ANASFV employs reference genomes to correct low-quality reads typical of Oxford Nanopore Technologies (ONT) sequencing, which guides the assembly process but may overlook open reading frames (ORFs) not present in the reference genomes. Additionally, the authors highlight that substantial host DNA contamination is a significant challenge in ONT sequencing, often leading to inadequate coverage. Rather than completely removing all reads that map to the host (*Sus scrofa*), this problem might be better addressed by removing only those reads that map exclusively to the host and not to the ASFV genome.

Regardless of the pipeline used, a suitable computational infrastructure and personnel with basic bioinformatics expertise are essential. However, we believe PePApipe may need lower computer resources than other pipelines as it was designed with the aim of providing simplicity in the first place, thus every step is clearly detailed, as can be found in the pipeline access websites.

## Methods

### DESIGN OF THE ALGORITHM

This pipeline, developed in Python, can be run using basic Linux commands to launch a Bash script with a Slurm protocol and can be executed on multi-sample batches. The work is sequentially performed over three main areas of genomic analysis: quality control and pre-processing of raw reads; *denovo* genome assembly and variant calling (Figure 1). Starting from raw data (paired .FASTQ files) obtained from short-read sequencing platforms (Illumina), ‘PePApipe’ implements in a sequential manner all crucial steps by using 13 software tools adequately built into a single pipeline.

**Figure 1.**
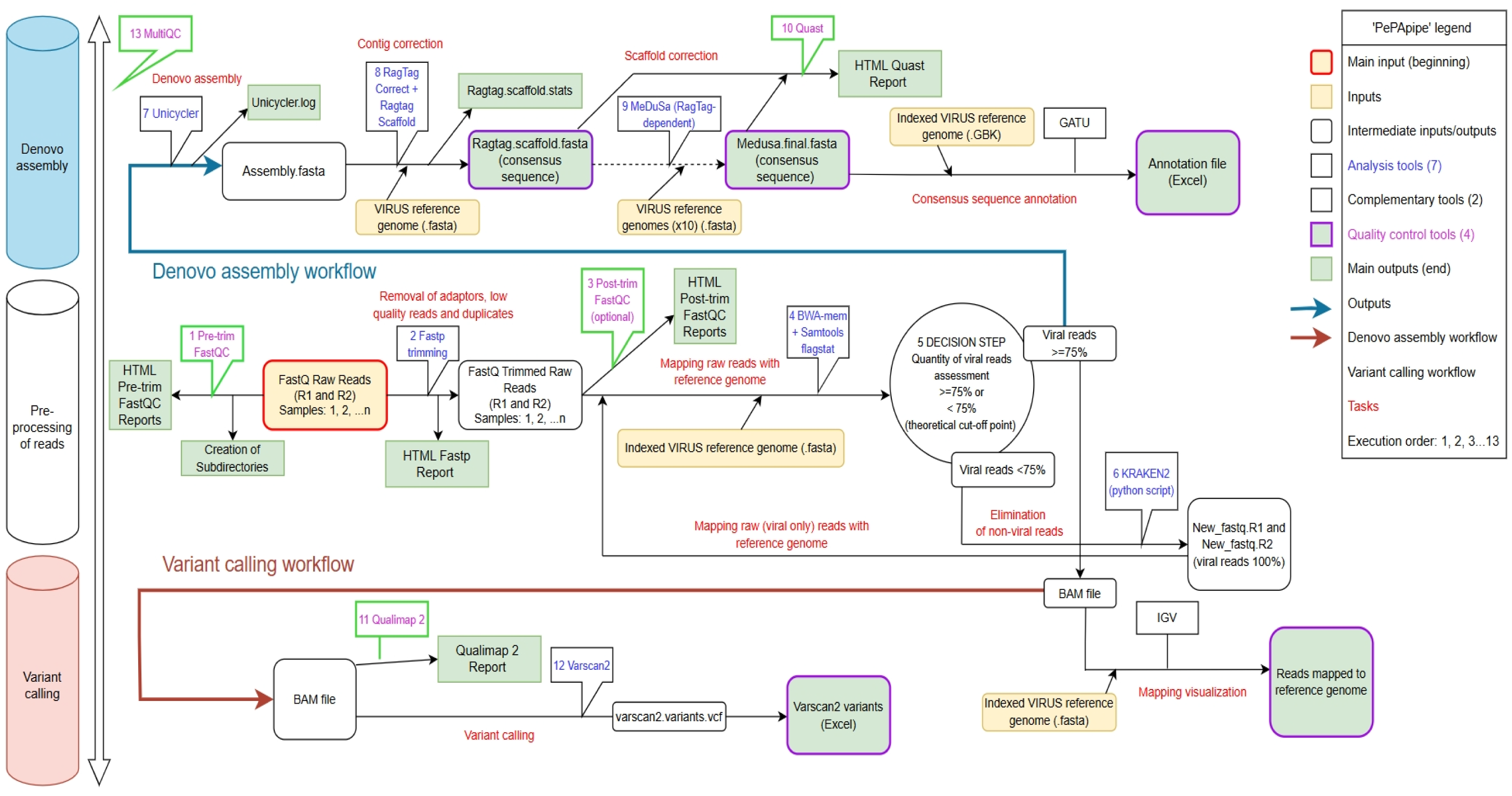
Overall bioinformatic analysis flow followed with PePApipe (in-house built Python pipeline specifically designed for ASF viruses).

**Figure 2.**
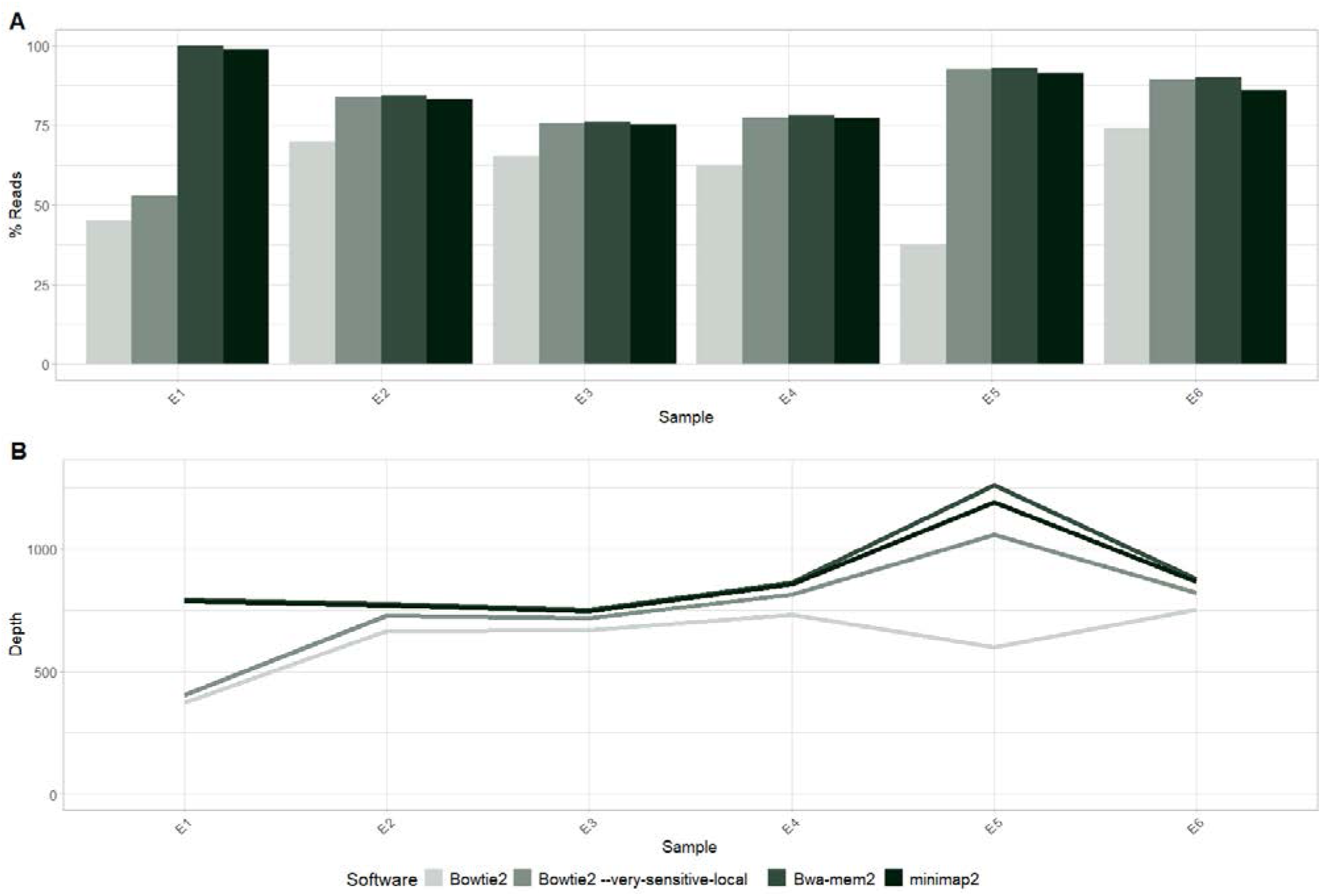
Comparison of Bowtie2, bwa-mem2 and minimap2 as mapping tools for ASFV sequences.

This pipeline can be run in a locally or remotely by accessing a computing server. A basic knowledge of Linux programming language is necessary both to install the programmes and to run the pipeline.

The pipeline with all associated information (including the pipeline itself and all accessory scripts) is accessible through the repositories Protocols.io (https://www.protocols.io/private/DE331EDEA81311F0AE550A58A9FEAC02) and GitHub (https://github.com/vilocha/PePApipe-pipeline) under the name ‘PePApipe: a complete bioinformatics analysis pipeline for African Swine Fever Virus genomes’.

### Detailed installation steps necessary to build the pipeline

The following steps must be implemented in the suggested order to successfully install all necessary programmes and be able to run this pipeline. The actual code lines used within the pipeline are clearly shown in the pipeline script itself accessible through GitHub and Protocols.io. These code lines are already built within the pipeline and will be executed at once when the pipeline is run.

The headers in the bash scripts specify the computation resources needed in the computer or server to allow the execution of the pipeline and its accessories.

According to needs, the different modules in the pipeline can be activated/ deactivated by just uncommenting/ commenting them in the main section of the pipeline script, respectively.

### SYSTEM AND METHODS IMPLEMENTATION

The input or starting point of the pipeline is always the set of raw reads which are the outputs of the sequencing platform (short reads from Illumina, although long reads from Oxford Nanopore Technologies can also be used) organized in two files: .FASTQ_R1, or forward reads, and .FASTQ_R2, or reverse reads. This is the standard output of pair-end sequencing protocols, which are common when working with Illumina protocols. The output files of the different steps in this pipeline are specified on the installation webpages accessible through GitHub and Protocols.io. We tested all steps with Illumina sequencing products (paired sampling) from six ASFV samples named E1 to E6 (38).

### STEPS

#### Quality control of reads

This is the very first step which is necessary to ensure a good quality of the .FASTQ files provided by the sequencing platform. The quality check of raw reads is carried out with the tool FastQC (13). As a general rule, the per-base sequence quality (Phred score) should always be ideally over 28, and it is recommended not to drop it below 20.

Once the quality control step is complete and one is satisfied that the quality of sequences is adequate to continue the processing, the reads are ‘cleaned’ with the tool fastp (14). This step is necessary to ensure that adapters and low quality and duplicate reads are removed off the .FASTQ files. In addition, fastp produces a post-trimming quality analysis report, making it unnecessary to perform a further stand-alone quality control with FastQC.

Only reads with a minimum Phred quality score of 20 (the default fastp Phred quality score is 15) and a minimum length of 15 base pairs (default parameter) are kept after applying fastp, but these parameters can be changed in the code line according to user needs. Depending on the number of reads available, these parameters can be changed up or down with the aim of maximizing output and minimizing low quality reads.

#### Viral reads content check

In this step the reads that passed the previous filters (‘clean reads’) are mapped against the virus reference genome in .FASTA format chosen for convenience in each particular case using the software tools BWA-mem2 (24) and Samtools (28).

This step is probably the most important of all steps included in this part of the flow, and of the pipeline for that matter. The viral genome reference used for convenience by the authors of the pipeline was ASFV Georgia 2007/1 (Genbank assembly: GCA_003047755.2), since it was the initial isolate of genotype II emerging in Eastern Europe in 2007 and is considered by the scientific community as the reference strain.

The objective of this mapping is two-fold: on the one hand it allows to quantify the amount of reads within the original .FASTQ files actually mapping with an ASFV (and thus to also quantify the amount of reads not mapping with an ASFV), and on the other hand, to create a set of reference-mapped contigs (.BAM file) as a first step in the search of genetic variants (SNPs and indels) in the genome of our virus problem compared to the genome of the virus reference used.

#### Quality check of reads mapped to reference genome of choice

One of the tools most commonly used for this purpose is Qualimap 2 (39). The rationale behind is to assess the quality of the mapping of trimmed raw reads to virus reference genome which was performed to calculate the amount of viral reads present. The outputs can be verified in .PDF and .HTML formats.

#### Quantitative assessment of viral reads (theoretical cut-off point: 75%)

It is important to carry out all bioinformatics analyses only on viral reads free of contamination with reads from other origins (host, environment or handling personnel) or, if this is not possible, at least on a number of viral reads as high as possible. This is because the results we obtain if the analyses are performed on a high number of non-viral reads may be spurious and therefore not representative of the virus we are trying to characterize.

This is a point in the pipeline where a decision has to be made as to the quality level we want to apply to our analysis in relation to the available reads (Figure 1). A theoretical threshold (cut-off point) of 75% has been established by the authors in order to decide whether to continue the analyses with the original .FASTQ files already filtered (‘clean reads’) or whether to carry out yet another filtering step to get rid of non-viral reads.

Apart from the methods used to isolate the viral DNA, the method used to prepare the libraries (with or without enrichment) is the single most influential step on the amount of foreign (non-viral) DNA reads found in our sample. If a non-targeted or non-enriched library preparation method is used, there will be a huge number of non-viral reads present in the .FASTQ files, thus these may be larger in size although not necessarily better in quality.

Hence, this decision threshold is arbitrary and may vary depending on the initial .FASTQ files and whether the library has been prepared using enrichment or not. Libraries prepared for shotgun sequencing are likely to contain <1% of viral fragments.

#### Removal of reads not belonging to Viruses

If we decide to carry out another filtering step to get rid of non-viral reads, a new set of .FASTQ files must be created by mapping the ‘clean reads’ against all published reliable reads belonging to the Superkingdom Viruses (NCBI: txid10239). To carry out the removal of non-viral reads a loop with an additional Python script using the tool Kraken2 (40) is necessary. The procedure consists of two steps: building of the virus database and extraction of the viral reads. This extra filtering step will create a new set of .FASTQ files containing only viral reads.

As already mentioned, this procedure can be performed whenever the chosen threshold of viral reads is not reached (in our case 75%) or it may also be systematically run every time, independently of the amount of non-viral reads present in the original ‘cleaned’ set of .FASTQ files.

If there is availability of a substantial amount of samples, a benchmark exercise can be carried out to refine the decision threshold or cut-off percentage point, making it lower or higher than 75% according to each specific situation.

#### *De novo* viral assembly

This step is mainly performed with the software tool Unicycler, and ultimately refined with the programmes RagTag (using Unicycler outputs as inputs) and MeDuSa (using RagTag outputs as inputs).

These tools use several dependency programmes in order to obtain a full viral *de novo* genome assembly starting from the .FASTQ files containing our already ‘cleaned’ reads, be these the original .FASTQ files or the new set created using Kraken2.

Although Unicycler (17) was originally built for bacteria, ideally combining both short (Illumina) and long reads (Oxford Nanopore Technologies), it performs equally well using only viral genomes short reads.

A most positive aspect about this type of assembly is that a viral reference genome is not used at all, thus no bias from this source can be introduced, allowing for a genuine assembly of the putative whole genome of our virus problem. This step is the most important within the *de novo* assembly part of the flow, but unfortunately it is also the most consuming in terms of time and resources.

There are particularities in relation to the three tools used in the *de novo* assembly. In order to obtain a proper assembly of our virus whole genome it is important to specify that what we are looking for is a linear sequence, in other words, that the expected number of linear (non-circular) sequences in the underlying sequence being assembled is just one, and so these requirements must be included in the code in the Unicycler section.

After the first assembly step with Unicycler is concluded, a correction of the obtained contigs converting these into scaffolds is performed with the RagTag tool (20). This step is necessary to guide the *de novo* assembly performed by Unicycler into plausible contigs belonging to the ASFV virus, using for that purpose the chosen viral reference genome in .FASTA format (in our case ASFV Georgia 2007/1; Genbank assembly: GCA_003047755.2).

It is important to note that, even though a viral reference is used, not a single fragment of it is incorporated into the assembly obtained with Unicycler, so the only activity performed in this step is that of contig correction and scaffolding. RagTag comprises four software tools (‘correct’, ‘scaffold’, ‘patch’ and ‘merge’) but only the first two are used in the pipeline. ‘Patch’ is not used to avoid adding viral reference fragments and ‘merge’ is not necessary since only ‘scaffold’ was used as scaffolding method.

After the correction and scaffolding performed by RagTag, a third tool called MeDuSa (21) is incorporated as part of the *de novo* assembly process. MeDuSa performs a scaffold correction in a similar way to RagTag, but instead of using only one viral reference it uses as many well curated references as we are able to provide, always making sure the references are akin to the ASFV genotype we are studying. Up to 10 references belonging to ASFV virus genotype II have been included in this pipeline, which in alphabetical order are: Arm_07_CBM_c2_LR812933.1, ASFV_Georgia_2007_1_FR682468.2.fasta, ASFV_HU_2018_MN715134.1, ASFV_LT14_1490_MK628478.fasta, Belgium_2018_1_LR536725.1, Estonia_2014_LS478113.1, Korea_YC1_2019_ON075797.1, POL_2015_Podlaskie_MH681419.1, Tanzania_Rukwa_2017_1_LR813622.1 and Ulyanovsk_19_WB_5699_MW306192.1.

The higher the number of references used, the better the result of this step will be. Briefly, the contigs obtained from the previous step are further assembled into larger scaffolds with MeDuSa. If the quality of the original reads is high this step generally results in one final contig, which is the putative consensus sequence of the virus under study. On the contrary, poor quality sequences may result in a large number of contigs, and if the number of contigs exceeds 3-5 it would be advisable to reject those sequences.

#### Quality assessment of the *de novo* assembly

This is a quality analysis that can be performed over all steps in the *de novo* assembly loop with the intention of investigating the overall quality of the process. The tool included in the pipeline for this task is Quast (QUality ASsessment Tool) (41) and the outputs are available on different formats: .TXT, .PDF, .TSV, .HTML, etc.

#### Search for viral genome variants (variant calling)

Several tools are used to reveal genetic variants (SNPs and indels) present in the virus genome under study when compared to a viral reference genome.

In general, there are two main types of tools: those that use a Bayesian inference and those that use a deterministic/numerical approach. The tool included in this pipeline is called VarScan 2 (30) and is one of the second type (numerical approach).

In order to obtain the most plausible record of variants with this tool the following five parameters have been defined with the specified values:

- Minimum number of reads with change to call a variant: 5
- Minimum depth (coverage=total number of reads supporting that variant): 20
- P-value: 0.001
- Minimum average quality: 30
- Minimum variant frequency in that position: 0.05 (to account for true minority variants)

We used a tool for variant calling provided by Qiagen CLC Genomics Workbench (31) to compare the variant calling results obtained with VarScan 2 to ensure that the latter performs adequately.

#### Overall quality assessment of the analysis

In order to perform an overall assessment of the analysis, the tool MultiQC (42) may be used. This tool evaluates all inputs and outputs in the working directory and performs a general quality summary assessment, which is based on the quality of the inputs (.FASTQ files) and on the type of programmes used. This tool must be executed in the working directory where all folders used during the running of the pipeline are located. This final step is valuable if we want to perform benchmark exercises to compare different inputs or different tools or both.

#### Expected outputs and their interpretation

The pipeline produces all necessary files to interpret the results, classifying them into folders, in an approximate time that depends on the size of the .FASTQ files, from 10 minutes with files of around 0.5 Mb through to 30 minutes with files of 1 Mb in size, or to longer times if high capacity sequencers are used. Several intermediate files are also created in the process, which can be used as inputs for further or parallel analyses, as well as to check for possible errors during the analysis. Among these complementary post-analysis steps are viral annotation with GATU (https://4virology.net/virology-ca-tools/gatu/) or genome visualization with IGV (https://igv.org/).

All steps are amenable to user control by means of switching on/off the necessary sections in each particular case. There are several steps where the quality of the processes can be checked to make sure the progress of the results is in line with the user expectations.

The pipeline can be easily adapted to viruses other than ASFV by changing the parameters relevant to the new virus and running the specific pipeline sections accordingly.

The most important outputs are the consensus sequence of the genome of the virus we are studying and the table of variants for that particular virus against the chosen reference genome. Secondary outputs are reference mapping parameters, *de novo* assembly parameters and quality assessment summaries.

PePApipe is user-friendly and can be installed through a straightforward series of steps. While it is primarily designed for analyses involving short reads, it is also compatible with long-read data. The outputs are consistently organized into folders, providing a structured basis for further analysis. Additionally, the code is open-source and can be modified and adapted to meet user specific needs, provided that appropriate credit is given.

## DATA AVAILABILITY

The primary data used in this paper is available at the ENA repository (https://www.ebi.ac.uk/ena/browser/home) under accession number PRJEB105327, from the 19^th^ December 2025.

## CODE AVAILABILITY

The code needed to run the analyses with all associated information (including the pipeline itself and all accessory scripts) is open source and accessible through the repositories Protocols.io (https://www.protocols.io/private/DE331EDEA81311F0AE550A58A9FEAC02) and GitHub (https://github.com/vilocha/PePApipe-pipeline) under the name ‘PePApipe: a complete bioinformatics analysis pipeline for African Swine Fever Virus genomes’. Support contact email for users is available at the websites above.

‘PePApipe’ is released under the MIT License.

Private Protocols.io link for reviewers: https://www.protocols.io/private/DE331EDEA81311F0AE550A58A9FEAC02

## FUTURE DIRECTIONS

The future plan is to include a viral genome annotation tool in the pipeline to harmonize this crucial and complex step for a large virus which can encompass more than 180 ORFs along the genome.

Finally, although the original aim of this study was to design a pipeline to specifically build and analyse the ASFV genome from NGS experiments, this can be easily adapted for the study and assembly of any other viral organisms.

## ACKNOWLEDGMENTS

Dr. Carmina Gallardo and Prof. Marisa Arias, from CISA-INIA-CSIC, are recognized for their contribution to provide the ASFV samples used for this study.

